# Spatial attention in perceptual decision making as revealed by response-locked classification image analysis

**DOI:** 10.1101/2022.07.30.499074

**Authors:** Hironobu Sano, Natsuki Ueno, Hironori Maruyama, Isamu Motoyoshi

## Abstract

In many situations, humans serially sample information from many locations in an image to make an appropriate decision about a visual target. Spatial attention plays a crucial role in this serial vision process. To investigate the effect of spatial attention in such dynamic decision making, we applied a classification image (CI) analysis locked to the observer’s reaction time (RT). We asked human observers to detect as rapidly as possible a target whose contrast gradually increased on the left or right side of dynamic noise, with the presentation of a spatial cue. The analysis revealed a spatiotemporally biphasic profile of the CI which peaked at ∼350 ms before the observer’s response. We found that a valid cue presented at the target location shortened the RT and increased the overall amplitude of the CI, especially when the cue appeared 500-1250 ms before the observer’s response. The results were quantitatively accounted for by a simple perceptual decision mechanism that accumulates the outputs of the spatiotemporal contrast detector, whose gain is increased by sustained attention to the cued location.

## Introduction

While humans and animals can immediately recognize scenes, objects, and materials at a glance [1, 2, 3, 4], in many situations they need to sequentially sample and integrate information to take more appropriate decisions about the world [5, 6]. In this serial vision process, attention (and eye movements) plays a critical role in the selection of relevant information from spatially distributed image inputs [5, 7].

A large body of psychophysical evidence suggests that focal attention on the target’s spatial location facilitates visual processing of the target [6, 8, 9, 10, 11, 12, 13, 14, 15]. In most of the studies, attention has been demonstrated to increase detection performance and accelerate the reaction to the target. However, such behavioral measures alone may be insufficient in explaining how attention affects the neural process of making decisions about the target, unless the results are compared across a variety of conditions.

In visual neuroscience, reverse correlation analysis has been widely applied to reveal information that determines the system response [16, 17]. This analysis has been applied not only to the responses of cortical neurons [18, 19, 20] but also to human behavioral responses for a variety of visual tasks [21, 22, 23, 24, 25, 26, 27, 28]. As a variant of the analysis, the classification image (CI) method allows the visualization of what information in the stimuli observers consider important for a given perceptual judgment [29]. In a typical experiment of the CI method, the observer’s responses to a visual target embedded in white noise are collected, and the information in the stimulus that affected the observer’s response is mapped out by analyzing the correlation between the noise and the response in each trial. The CI method has been widely used to reveal the spatiotemporal distribution of critical information (or the perceptive field) that determines the observers’ judgments for various visual tasks with static and dynamic stimuli [30, 31, 32, 33, 34, 35].

Eckstein et al. (2002) applied the CI method to Posner’s cueing paradigm [8] and showed that the weight of information in the CI is greater at the spatial location where attention was directed [31]. However, observers made judgments after the visual stimuli had been shown, like in many psychophysical reverse-correlation studies [23, 24, 25, 27, 28]. Such post-stimulus judgment, usually based on the visual working memory, is not necessarily representative of the on-the-fly judgments that we make in real life. It is desirable to examine the effect of dynamic attention to the CI in the period before the observer responds to the shown target.

To clarify when observers make decisions during observation and what information they rely on to make decisions during observation, we can analyze correlations at each time point of the stimulus locked to the reaction time (RT) of the observer during the presentation of the stimulus rather than at the stimulus onset. Several studies have adopted this response-locked reverse correlation analysis in investigating the dynamics of perceptual decision making [22, 36, 37, 38]. Maruyama et al. (2021) recently applied response-locked CI analysis to a very basic visual task, namely luminance contrast detection [39]. Using a visual display similar to that used by Neri & Heeger (2002) [32], they measured responses and reaction times for a target stimulus slowly emerging from dynamic noise and calculated the correlation between the noise and response at each time point, locked to the observer’s reaction time (RT), in reverse chronological order. Adopting this protocol, they examined what signals and what point in the stimulus determined the observer’s decision about the target and RT, and they revealed spatiotemporally biphasic CIs and their RT-dependent variability. They further suggested that the results could be quantitatively reproduced by a simple computational model incorporating a perceptual process approximated by a spatiotemporal filter and a (drift-diffusion) decision process that accumulates the output.

In the present study, we extended the above experimental protocol to investigate the effects of spatial attention. To this end, in addition to noise and target stimuli similar to those used by Maruyama et al. (2021) [39], we presented a cue indicating the target’s location (a valid cue) or another location (an invalid cue) as used in Posner’s paradigm. Using this display, we measured responses and reaction times (RTs) for a target stimulus that appeared gradually in dynamic noise, and we compared RTs and CIs between the valid and invalid cue conditions. We found that the overall amplitude of temporally biphasic CIs was larger for the valid cue than for the invalid cue in the temporal period that the RT was shortened by attention. These results were well explained by a simple perceptual-decision model [39], with the introduction of a single additional assumption that spatial attention sustainedly increases the gain of the perceptual process.

## Methods

### Observers

We initially recruited seven participants, which was the same number as in the previous study [39]. However, the data of one observer whose reaction time was found to be exceptionally long were excluded in the early phase of experiment. The results of the others showed that one of the expected effects – an interaction between the cue condition and the peak of the amplitude – was statistically significant. Based on the effect size in this result, we conducted an a priori power analysis using G*Power 3.1 which revealed that a sample size of ten should be sufficient to achieve .95 power [40]. We then recruited four more participants, and finally ten human observers with corrected-to-normal vision, including three of the authors and seven naïve paid volunteers (having an average age of 24.2 years), participated in the experiment. All experiments were conducted in accordance with the Declaration of Helsinki. The study was approved by the Ethics Committee of The University of Tokyo. All observers gave written informed consent.

### Apparatus

Visual stimuli were generated on a personal computer and displayed on liquid-crystal display monitors. To deal with the COVID-19 situation, the monitors (three BENQ XL2720B, three BENQ XL2730Z, one BENQ XL2735B, one BENQ XL2430T, one BENQ XL 2731K, one SONY PVM 2541A, and one SONY PVM-A250) were set up in the participants’ own homes. The mean luminance of uniform backgrounds ranged from 44 to 115 cd/m^2^. All monitors had gamma-corrected luminance as calibrated with a colorimeter (ColorCal II CRS). The frame rate was 60 Hz. The viewing distance was adjusted to achieve a pixel resolution of 0.018 deg/pixel.

### Stimuli

The visual display followed that of the previous study [39]. The stimulus was a pair of square images of dynamic one-dimensional noise (4.6 × 4.6 deg), each comprising 16 vertical bars with a width of 0.29 deg (Fig. 1). The contrast (*C*_*noise*_ (*t*)) of each bar switched at a frame rate of 30 Hz according to a Gaussian distribution having a standard deviation of 0.1. The total stimulus duration was 4500 ms. Two independent one-dimensional noise fields were presented adjacent to the left and right of the gaze point.

**Figure 1.**
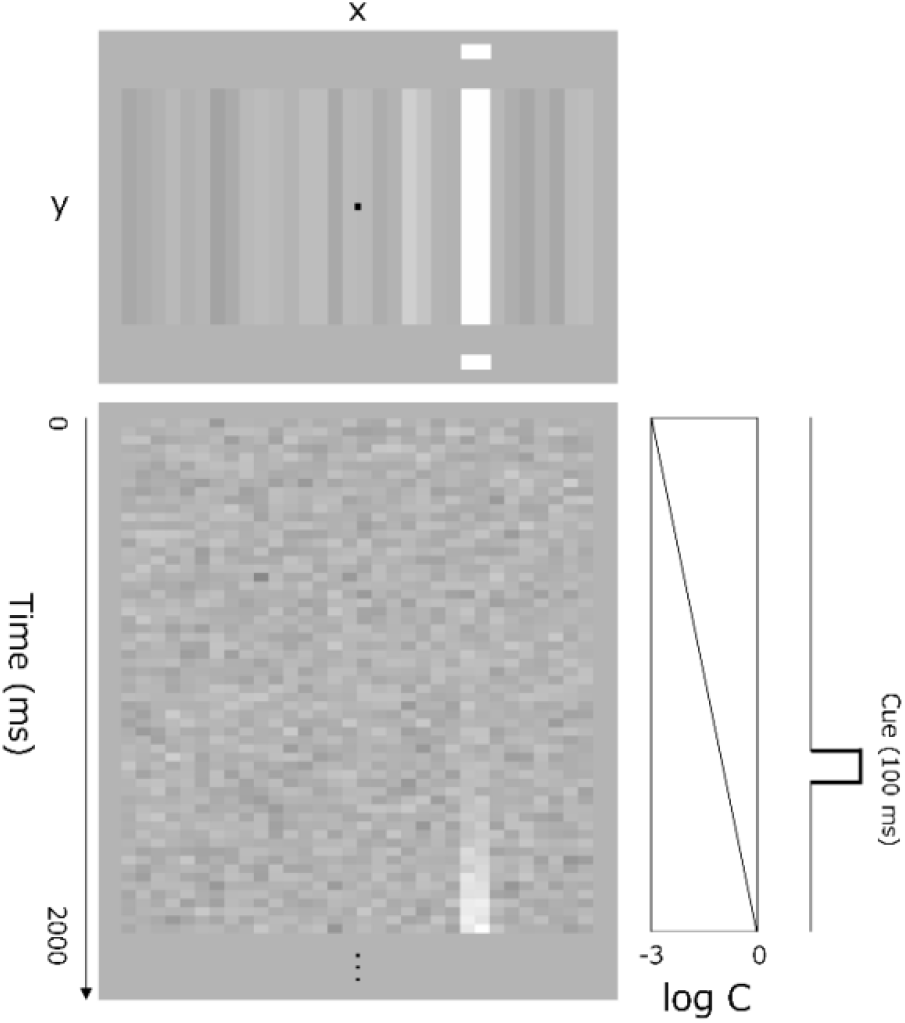
Schematic diagram of the visual stimuli used in the experiment. The upper panel shows a snapshot of a single stimulus frame. The bright bar on the right is the target, and the small horizontal bars above and below are the cue. The lower panel shows an X-T plot of the luminance variation of each bar. The target appears slowly in the right field. As shown in the right plot, the cue was presented for 100 ms with random timing from 500 to 2000 ms following the beginning of the increase in target contrast.

The target signal (*C*_*target*_ (*t*)) was added linearly to the two bars in the center of the noise area on either side. The addition of the target signal was set to start at a random frame between 0 and 500 ms after the start of the noise presentation.

The luminance contrast of the target stimulus, *C*_*target*_ (*t*), increased linearly on a logarithmic scale with time (*t*) according to

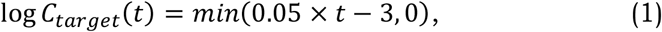

where *t* is the frame number (33 ms per unit) from stimulus onset. The contrast of each bar was clipped in the range from −1 to +1. The two fields, with and without the target signal, were transformed into a luminance image using the relation L(*t*) = L_mean_ (1 + C(*t*)), where L_mean_ is the mean luminance of the uniform background, which depended on the monitor of each observer (44–115 cd/m^2^).

In addition to the noise and target, a cue that probabilistically indicated the location of the target was presented. The cue was a pair of bright rectangles having a height of 0.3 deg and width of 0.6 deg. They were flashed for 100 ms at 0.6 deg above and below the central two bars in either the left or right field. The cue was presented at random timings of 500–2000 ms after the beginning of the increase in target contrast. The cue was presented in 96% of the trials. In 75% of these trials, the cue was presented at the target location (i.e., the valid-cue condition), and in the remaining 25% of the trials, it was presented on the opposite side (i.e., the invalid cue condition).

### Procedure

Observers binocularly viewed the display with a steady gaze at the fixation point and were asked to indicate by pressing a button whether the target appeared in the left or right field as rapidly as possible. Observers were informed that the cue would be presented at a valid location in the majority of trials. Auditory feedback was given for errors and responses that exceeded the deadline (4500 ms), and data in those trials were excluded from the analysis. The average error rate was 2%. The next trial started no less than 0.5 s after the observer’s response. Each session of the experiment comprised 160 trials, and sessions were repeated until at least 3200 trials (4960 at maximum) were completed for each observer. In the analysis, all trials in which the observer responded before the cue, regardless of the validity of the cue, were included in the no-cue condition, along with trials in which the cue was not presented until the end.

## Results

### Reaction time

Figure 2a shows the reaction time (RT) as a function of the cue onset time. Each data point is the mean RT in each 100-ms epoch from the cue onset. The RT of the individual observer is defined as the harmonic mean across trials in each epoch. The red lines show the results for the valid cue, the blue line those for the invalid cue, and the black line those for no cue. RT is ∼100 ms shorter for the valid cue than for the invalid cue, suggesting the effect of cueing. It is seen that the facilitation is weakened and diminishes when the cue is presented at ∼1500 ms or later, but this is simply because this range contained slow-response trials in which observers responded after the contrast of the target had become high. A two-factor ANOVA revealed significant main effects of the cue condition (F(2,18) = 48.48, *p* < 0.0001, η^2^ = 0.1011) and cue onset time (F(14,126) = 33.37, *p* < 0.0001, η^2^ = 0.0761) and a significant interaction between them (F(28,252) = 21.18, *p* < 0.0001, η^2^ = 0.0660).

**Figure 2.**
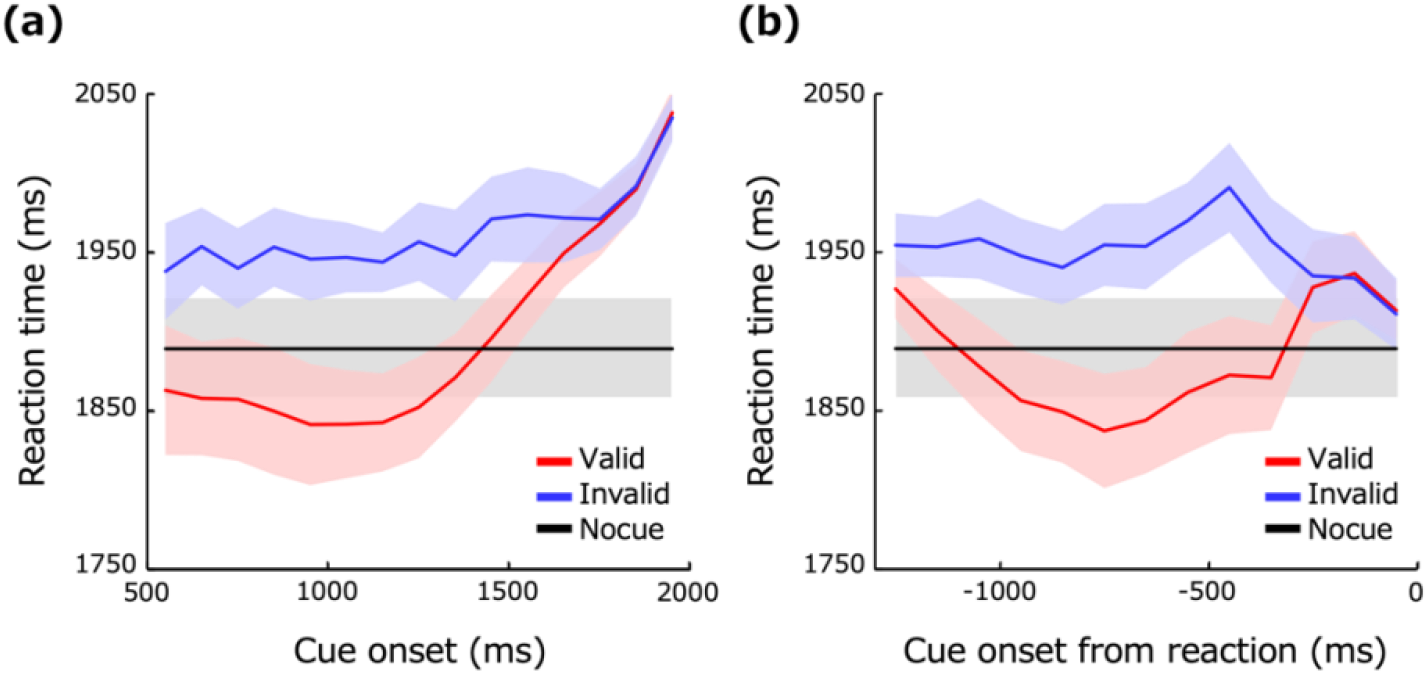
Effects of the spatial cue on the reaction time. (a) Reaction time as a function of the timing of the cue onset. Data are for each 100-ms epoch. The vertical axis shows the RT for the valid cue condition (red line), invalid cue condition (blue line), and no-cue condition (black line). The light-colored bands represent ±1 s.e.m. across observers. (b) Reaction time as a function of the time from the response to cue onset. Data are for each 100-ms epoch.

To better capture the dynamic variation in RT due to spatial cueing, we also calculated the mean RT as a function of the timing of the cue onset from the observer’s response as shown in Fig. 2b. The plot shows that the RT was especially shortened when the valid cue was onset 500–1000 ms before the response. The RT was particularly long when the invalid cue was onset 500 ms before the response. We also find that the RT become longer when the cue was onset 1000 ms before the response, but this is simply because most of the responses in these trials were exceptionally slow regardless of the validity of the cue. It may appear strange that the difference in the RT between the valid and invalid cue conditions was small when the cue was onset ∼300 ms before the response, but this difference was small simply because the cue was presented during the motor delay between the decision and button press (see also the section on computational modeling). A two-factor ANOVA revealed significant main effects of the cue condition (F(2,18) = 52.36, *p* < 0.0001, η^2^ = 0.1064) and cue onset from the response (F(14,126) = 15.14, *p* < 0.0001, η^2^ = 0.0499) and a significant interaction between them (F(28,252) = 18.81, *p* < 0.0001, η^2^ = 0.0602).

### Classification image analysis

Using the same procedure as in the previous study [39], we conducted reverse correlation analysis of the contrast of each bar and the observer’s response (left, right) at each time point (*t*) back from the reaction time, and we calculated the Classification Image (CI). Fig. 3a is a diagram of the analysis. As in the work of Eckstein et al. (2002) [31], CIs were calculated separately for the noise field where the target was presented and that where the target was not presented.

**Figure 3.**
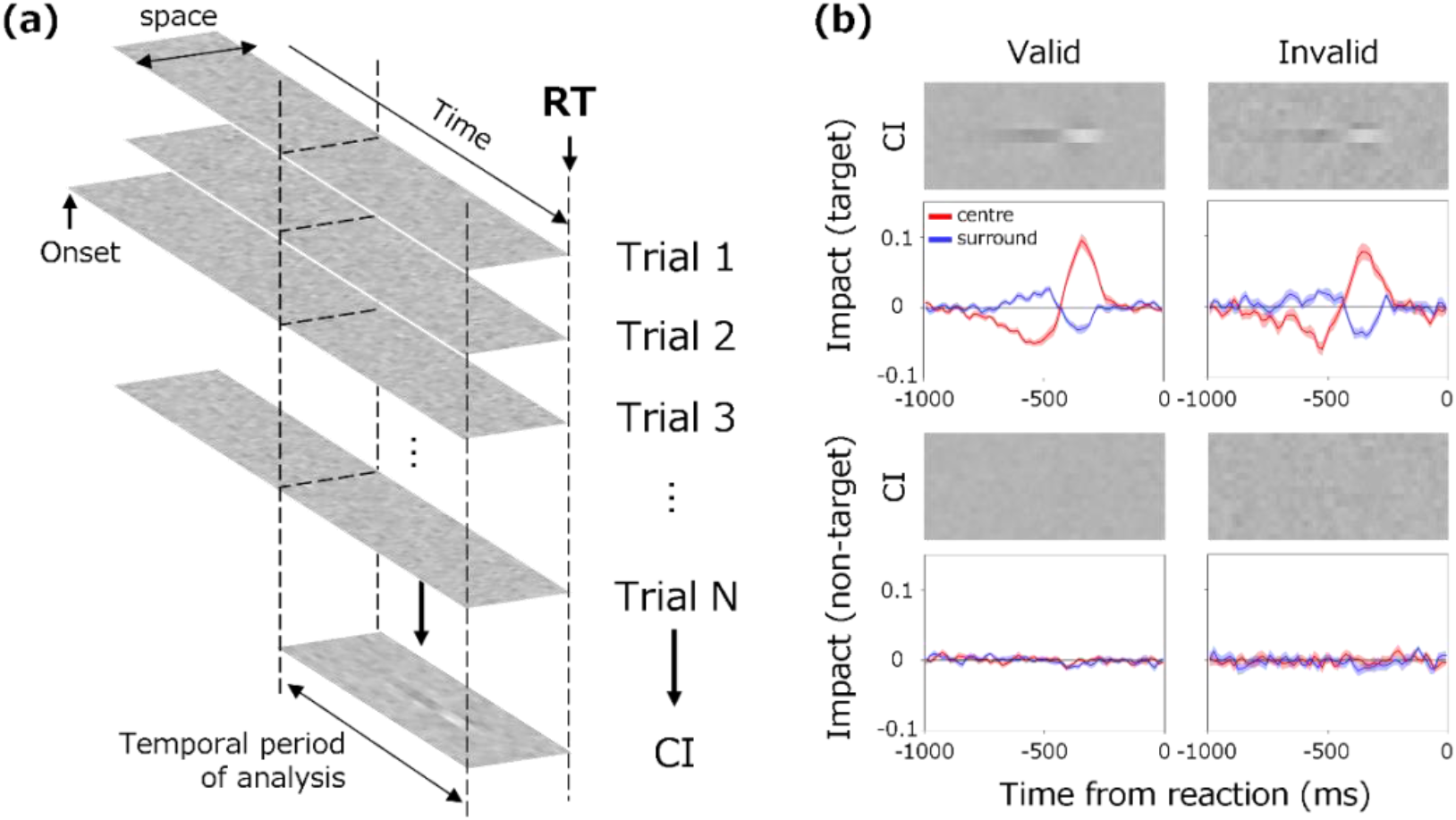
(a) Diagram of the response-locked classification image. The classification image (CI) was calculated for each bar contrast at each spatiotemporal position backward from the observer’s response time. (b) The upper images show CIs for the valid (left) and invalid (right) conditions, where the vertical axis indicates space and the horizontal axis indicates the time from the response. The lower curves show the average weights of the two bars in the center (red curves) and the average weights of the two adjacent bars (blue curves). The light-colored bands represent ±1 s.e.m. across observers. The upper panels show the results for the target field and the lower panels show the results for the non-target field.

The images in Fig. 3b show the average CIs obtained for the target (upper panels) and non-target (lower panels) fields. For each, the left and right panels respectively show the results for valid and invalid cue conditions. The brightness of each pixel represents the weight of noise on the response at spatial each position (vertical axis) and at each time before the response (horizontal axis). Bright pixels represent positive weights and dark pixels represent negative weights. The curves below the CI show the average weights of the two central bars (red) and the two adjacent bars (blue) in the CI. We refer to these as impact curves.

The CIs and impact curves obtained for the two cue conditions have similar trends, except that the amplitude appears slightly larger under the valid cue condition. For both cue conditions, there is clearly a characteristic spatiotemporal biphasic profile immediately before the response in the target field (upper panels). At the center of the CI where the target appeared, positive weights peak at approximately 350 ms before the response and negative weights at approximately 500 ms before the response. The opposite profile is found on both sides surrounding the center. These results, having clear agreement with the results of the previous study [39], suggest that the observer’s decision is triggered by the luminance increase after the luminance decrease in the central bars and the relative emphasis of these luminance variations in space. Meanwhile, CIs of the non-target field (lower panels) show no systematic variation, and we will not consider these data in the following analysis.

The RT data shown in Fig. 2b indicate that the observer’s response was systematically accelerated when the valid cue was presented at a particular time before the response. With reference to these results on the RT, we next examine whether the CI varies between the valid and invalid cue conditions depending on how long before the response the cue appears. To this end, we divided the trials into epochs of 500 ms each based on the cue onset time from the response and calculated the CI for each epoch with a shift of 250 ms.

Figure 4a shows the CIs and impact curves for different times from the response to cue onset in the target field. The impact curves preserve a biphasic shape regardless of the cue onset time backward from the response. There is a negative weight of the center impact curve when the cue onset was long before the response. Given that trials in which the cue appeared long before the response are dominated by long RTs, this reflects the fact reported in the previous study that the negative weight is larger in trials of longer RTs [39].

**Figure 4.**
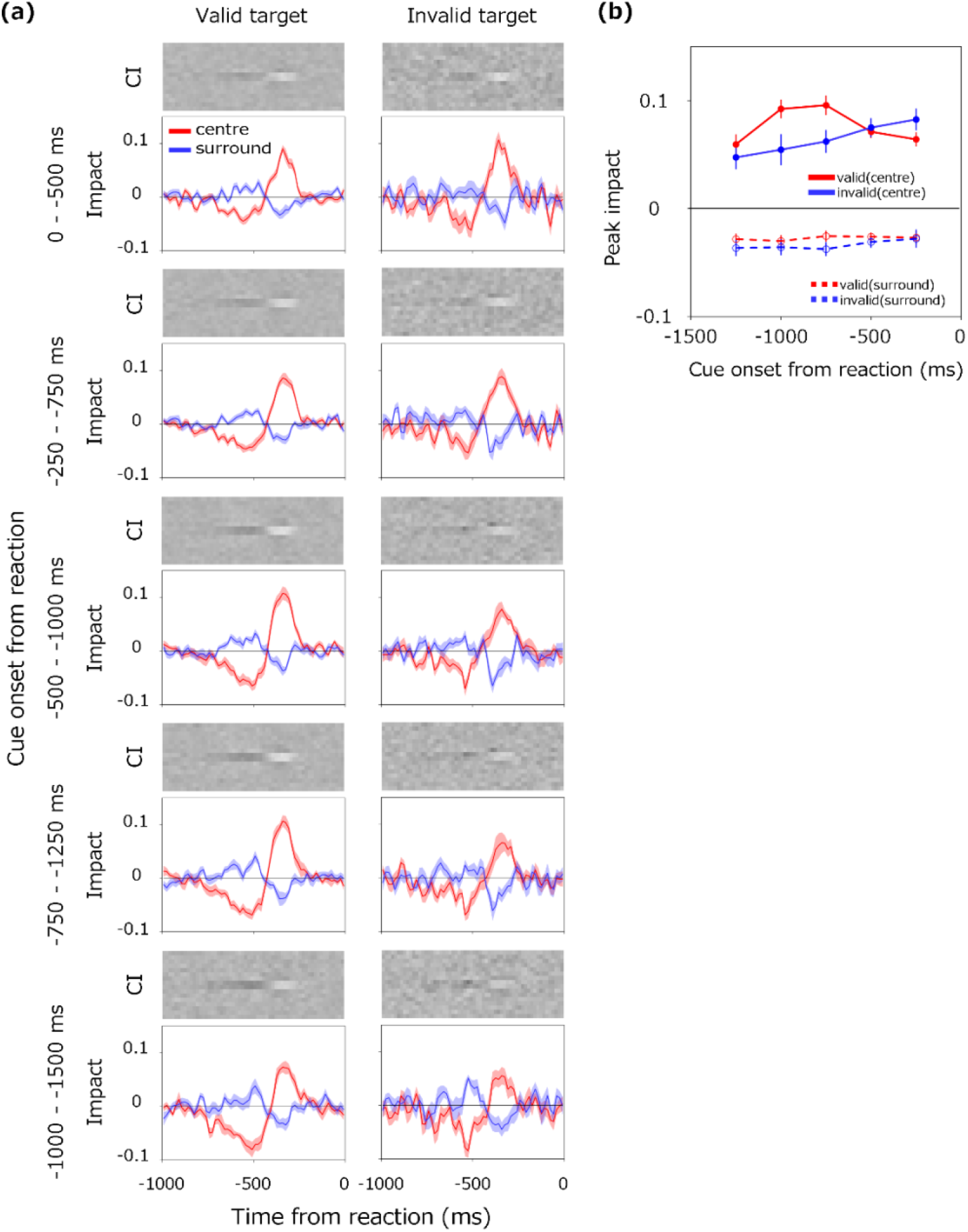
(a) CIs and impact curves calculated for different times from the response to cue onset in the target field. From the upper to the lower panels, the results are for trials with the cue appearing shortly to long before the reaction. The left panels show the results for the valid cue condition and the right panels the results for the invalid cue condition. (b) Peak impact at approximately 350 ms before the response plotted as a function of the time from the response to cue onset. Red and blue circles represent results for the valid and invalid cue conditions, respectively. The solid and dashed lines show the results for the center and surrounds, respectively. Error bars are ±1 s.e.m. across 10 observers.

The overall amplitude of the impact curve is visibly different between the two cue conditions at a specific cue onset time backward from the response. Fig. 4b plots the peak of the impact curve for each epoch. Here, the peak of the amplitude is defined as the average of the impacts within the range of 300–400 ms backward from the response. The red circles represent results for the valid cue condition and the blue circles those for the invalid cue condition. The solid line presents the results for the center impact and the dashed line those for the surrounds impact. We find that the positive weight is considerably larger for the valid cue condition than for the invalid cue condition when the cue was onset 500–1250 ms before the response, with the RT being especially shortened by the valid cue. This can be interpreted as the spatial attention being guided by the cue boosting the impact of information at that location.

When the cue was onset 0–500 ms before the response, we find that the difference between the two cue conditions is small, or the weights under the invalid cue condition appear to be slightly larger. This effect may appear to reflect suppressive effects of the cue onset itself on the response, but it is more likely a result of inclusion of trials in which the decision was made before the cue was presented, as was the case for the RT data (Fig. 2b).

A two-factor ANOVA was performed on the data in Fig. 4b for the cue condition (valid, invalid) and the time from the response to the cue onset (five epochs). The results show that the main effect of the cue condition (F(1,9) = 4.40, *p* = 0.0654, η^2^ = 0.0343) was not significant, but the main effect of the time from the response to the cue onset (F(4,36) = 4.80, *p* = 0.0033, η^2^ = 0.0712) and the interaction (F(4,36) = 8.80, *p* < 0.0001, η^2^ = 0.1060) were significant. Corresponding t-tests conducted for each epoch showed significant differences for 0–500 ms (t(9) = −2.59, *p* = 0.0291, d = 0.8198), 500–1000 ms (t(9) = 3.38, *p* = 0.0082, d = 1.0671), and 750–1250 ms (t(9) = 3.04, *p* = 0.0141, d = 0.9598). ANOVA on the peak impact for the surrounds showed no significant effect.

## Discussion

The present study applied the response-locked CI method to analyze the effect of attentional cueing on dynamic decision making in a simple contrast detection task. Consistent with the results of many previous studies [8, 9, 10, 41], we found that the RT was shortened by the valid cue, especially in the period of 300– 1100 ms from cue onset. Furthermore, although the CIs had a spatiotemporally biphasic profile as in the previous study [39], their overall amplitude was boosted by the valid cue, which also shortened the RT.

The reduction of the RT by the valid cue began to appear 300 ms from cue onset and was long lasting. Importantly, this response facilitation was observed even when the valid cue was presented more than 500 ms before the response. These results suggest that the valid cue summoned the observer’s sustained attention to the target location and facilitated the response.

When cue the was onset 500–1250 ms before the response, CIs showed clear differences in the overall amplitude of the CI between the two cue conditions, while their spatiotemporal tuning profiles were relatively constant. Under the valid cue condition, the cue directed attention to the target region and increased the weight of information, thereby enhancing contrast detection sensitivity. Under the invalid cue condition, the cue directed attention to the opposite side of the target, which relatively de-emphasized information in the target region and suppressed contrast detection. These results are consistent with a large body of psychophysical evidence indicating an increase in the contrast sensitivity to stimuli at an attended location [13, 14, 15, 42, 43, 44].

It is noteworthy that the variability of the RT due to a spatial cue was ∼100 ms or more. This variability is considerably larger than values reported in many attention studies [8, 9, 41]. If we convert this 100-ms difference to the target bar contrast, we can infer that the valid cue enabled observers to detect targets with nearly half (∼0.3 log units) the contrast of targets detectable with the invalid cue. This substantial increase in behavioral sensitivity seems to be due to the fact that we employed targets with gradually increasing contrast over time. Psychophysical evidence indicates that attention increases contrast sensitivity by 0.05–0.1 log units for targets of a stepwise waveform [13, 14] but by as much as 0.5–1.0 log units for targets with a gradual onset [44]. This large sensitization for the gradual target is observed even in the absence of background noise, indicating an increase in the absolute detection sensitivity (i.e., reduction of internal noise) rather than the reduction of external noise [44].

### Computational model

On the basis of the above considerations, we examined whether the present results can be accounted for by a standard model of perceptual decision making with an additional assumption of the attentional enhancement of the perceptual sensitivity. The previous study assumed a hybrid model comprising a decision process that serially accumulates the outputs from the perceptual process, and it successfully predicted RTs and CIs in a similar display without spatial cueing [39]. Here, we extended the model by incorporating the effect of attention as an increase in the gain of the perceptual process, and we attempted to predict the characteristic variability of RTs and CIs depending on the cue validity.

Figure 5 presents an outline of the model. The model compares the spatially summarized outputs between two fields of the perceptual process approximated using a linear spatiotemporal filter. The decision process accumulates the differential signal of the two fields as sensory evidence over time and makes a decision when the evidence reaches a given boundary. The perceptual process is approximated as a spatiotemporal linear filter, and the decision process follows the standard drift-diffusion model for a two-alternative forced-choice task [45, 46, 47, 48, 49, 50]. Fig. 5 specifically illustrates each step in the case that the target appears on the left. In the model, the effect of spatial attention at the cued location is implemented as an amplification of the perceptual response in either the left or right field. The computation of each step is described in detail below.

**Figure 5.**
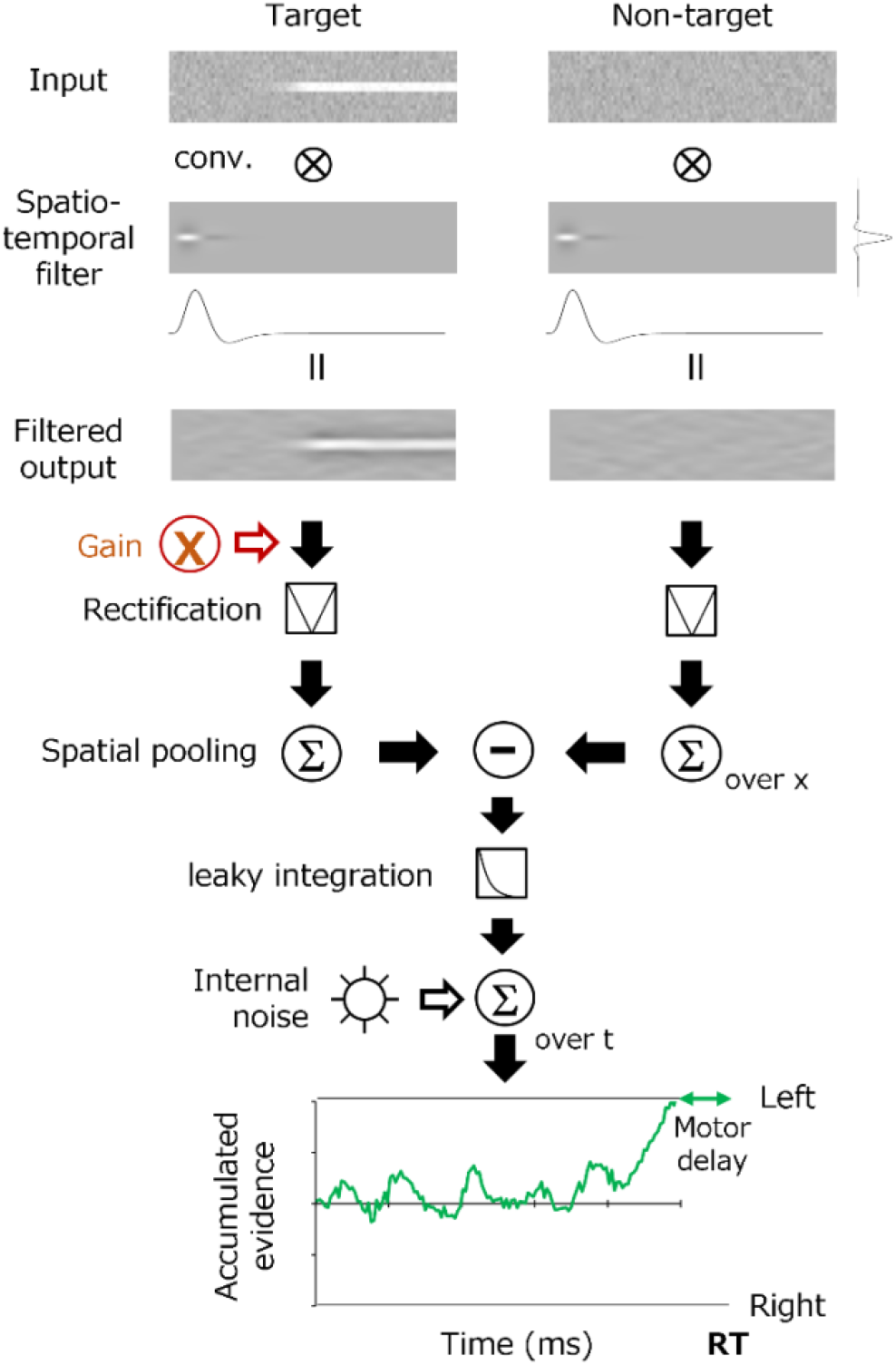
Schematic diagram of a model based on spatiotemporal filtering and the accumulation of sensory evidence. See the main text for details.

The perceptual system is approximated as a space–time separable linear filter, *F*_*st*_(*x, t*), written as

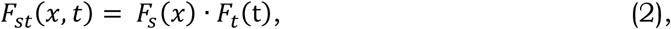

where *F*_*s*_ (*x*) is the spatial filter, *F*_*t*_(t) is the temporal filter, *t* is the frame number (33 ms per unit), and *x* is the pixel (0.04 deg per unit). The spatial filter *F*_*s*_(*x*) is given as a difference-of-Gaussians function, which has been widely used as a first-order approximation for contrast detectors in the early visual system [51]:

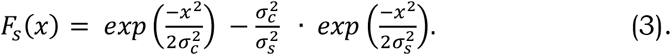

Here, *σ*_*c*_ is the standard deviation of the center and *σ*_*s*_ is the standard deviation of the surrounds. The temporal filter *F*_*t*_(t) is given as a biphasic function [52, 53]:

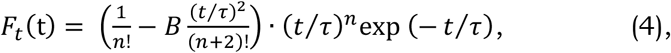

where *n* is the number of stages of the leaky temporal integrator, *τ* is the transient factor, and *B* is a parameter that defines the amplitude ratio between the positive and negative weights. The response of the perceptual system *R*(*x, t*) is then obtained by convolving the spatiotemporal filter *F*_*st*_ (*x, t*) with the stimulus input *I*(*x, t*). Here, *A* is the gain of the system. Note that the gain *A* is assumed to be 1 typically and amplified in fields to which attention is directed by spatial cuing.

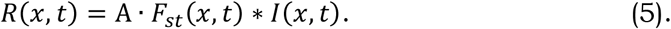

Determination of whether the target is presented in the left or right field is made by comparing the spatial sum of the absolute values of the responses in each field. The modeling is assumed to continuously monitor the difference ΔR(*t*) between the left and right responses at time *t* from the stimulus onset. Here, ΔR(*t*) is considered the sensory evidence at time *t* in the decision process:

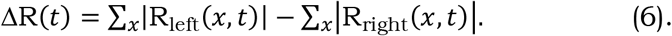

Decisions abouts targets are based on evidence accumulated over time. However, many decision-making studies suggest that sensory evidence decays with time [24, 28, 54]. This process is approximated by leaky temporal integration, which is mechanically ascribed to adaptive gain control [55, 56]. The model thus assumes the cumulative evidence *S*(*T*) at time *T* is obtained from an approximation of the noisy leaky integral of ΔR(*t*) as

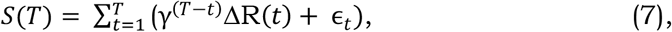

where *γ* is the time constant of evidence integration and ϵ_*t*_ is the internal noise having a normal distribution. The model observer makes a decision about whether the target is on the left or right when *S*(*T*) reaches a certain decision boundary; i.e., *b* or −*b*, respectively. Finally, the model observer is assumed to execute a manual response after a constant motion delay of 250 ms from *T*.

Our psychophysical data suggest a reduction in RT and an increase in the CI amplitude for the target at the cued location. According to the model architecture described above, this attentional facilitation must occur prior to the comparison of the left and right fields. We therefore chose to incorporate the effect of attention as a sustained increase in the gain of the perceptual system following the cue onset. Specifically, we assumed that the increase in gain due to attention begins with a fixed delay of 50 ms following the cue onset.

In our model simulations, we attempted to reproduce the impact curve (Fig. 4) for the data of each observer, using the stimuli actually shown to the observer. The parameters of the model were estimated to minimize the RMS error with the impact curve, separately for each of the valid and invalid cue conditions. Under the invalid cue condition, the gain (*A*) was fixed at 1. In accordance with previous studies [39], the number of stages of the temporal filter (*n*) was fixed at five.

Figure 6 shows the results of model simulation. It is seen that the model reproduces many aspects of the human data. Fig. 6a shows a continuous reduction in the RT under the valid cue condition. Figs. 6b and c presents larger amplitudes of the CI under the valid cue condition than under the invalid cue condition, except for the period immediately after the cue onset.

**Figure 6.**
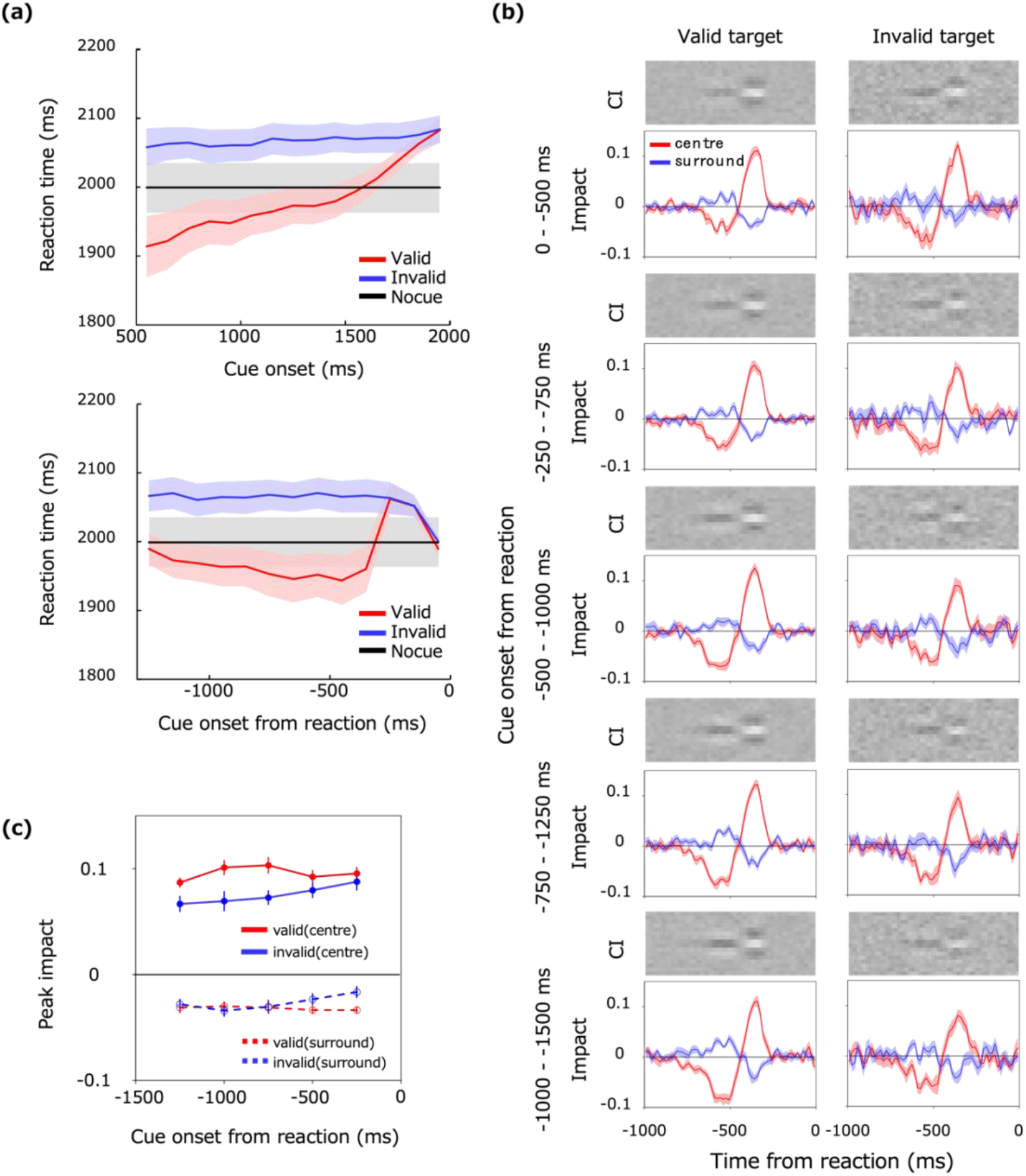
Reaction times and CIs predicted by the model. Each panel corresponds to the results for human observers in Figs. 2 and 4. (a) Variation in the RT with cue. RTs for the no cue condition were estimated by averaging the valid and invalid conditions. (b) CIs and impact curves for each 500-ms epoch of the cue onset from the response. (c) Peak impact near 350 ms before the response.

Estimated parameters and the s.e. across model observers were [*σ*_*c*_, *σ*_*s*_, *B, τ, γ, b*, ϵ_*t*_, *A*] = [2.94, 16.9, 0.505, 2.38, 0.207, 356.0, 10.6, 1.20 (s.e. = 0.13, 0.93, 0.006, 0.11, 0.004, 7.09, 0.69, 0.025)] for the valid cue condition and [*σ*_*c*_, *σ*_*s*_, *B, τ, γ, b*, ϵ_*t*_] = [2.92, 15.7, 0.490, 2.41, 0.212, 353.5, 9.80 (s.e. = 0.13, 0.59, 0.006, 0.10, 0.007, 7.22, 1.40)] for the invalid cue condition. The relative gain *A* of the spatiotemporal filter (1.20) is obviously greater than 1 (t(9) = 7.69, *p* < 0.0001, d = 2.431) for the valid cue condition, providing evidence of the sensory amplification by attention. Meanwhile, the other parameters did not significantly differ between the valid and invalid cue conditions, except for the ratio of positive to negative phases (*B*) in the temporal filter, which was significantly larger under the valid cue condition (t(9) = 2.47, *p* = 0.0355, d = 0.7816), but the difference was very small (0.505 vs. 0.490).

Taken together with the results of the previous study [39], the present results suggest that perceptual decision making in simple contrast detection can be well described by a simple perceptual decision model. Adding to this basic result, the present data suggest that the facilitation of behavioral decision making via spatial attention is explained by the increased gain of the perceptual process.

However, it is noted that the present finding does not rule out a variety of other possible explanations. Even assuming the model architecture shown in Figs. 5, it is difficult to determine exactly at what stage the attentional modulation of gain occurs. Our model assumes that the sensitivity of the spatiotemporal filter is modulated, but it is also possible that the spatially summarized output of the perceptual processes is modulated. Furthermore, if we assume another type of decision model such as a ballistic model [57, 58], which can deal with two sensory evidence signals independently for each of the two target fields, one cannot discriminate whether attention affects the perception process for each target or the decision process for each evidence signal. As pointed out in previous studies [38], it is generally difficult to distinguish between the involvements of perceptual and decision processes in the data of psychophysical reverse-correlation.

## Acknowledgements

This study was supported by JSPS KAKENHI 20H01782.

## References

1. Thorpe, S., Fize, D. & Marlot, C. Speed of processing in the human visual system. Nature 381(6582), 520–522 (1996).

2. Oliva, A. & Torralba, A. Building the gist of a scene: The role of global image features in recognition. Prog. Brain Res. 155, 23–36 (2006).

3. Motoyoshi, I., Nishida, S. Y., Sharan, L. & Adelson, E. H. Image statistics and the perception of surface qualities. Nature 447(7141), 206–209 (2007).

4. Whitney, D. & Yamanashi Leib, A. Ensemble perception. Annu. Rev. Psychol. 69, 105–129 (2018).

5. Treisman, A. M. & Gelade, G. A feature-integration theory of attention. Cogn. Psychol. 12(1), 97–136 (1980).

6. Wolfe, J. M. Visual search in The Handbook of Attention, 27–56 (2015).

7. Duncan, J. Selective attention and the organization of visual information. J. Exp. Psychol. Gen. 113(4), 501 (1984).

8. Posner, M. I. Orienting of attention. Q. J. Exp. Psychol. 32, 3–25 (1980).

9. Posner, M. I. & Cohen, Y. Components of visual orienting in Attention and performance X: Control of Language Processes 531–556 (Hillside, NJ: Erlbaum, 1984).

10. Nakayama, K. & Mackeben, M. Sustained and transient components of focal visual attention. Vis. Res. 29(11), 1631–1647 (1989).

11. Lee, D. K., Koch, C. & Braun, J. Spatial vision thresholds in the near absence of attention. Vis. Res. 37, 2409–2418 (1997).

12. Lee, D. K., Itti, L., Koch, C. & Braun, J. Attention activates winner-take-all competition among visual filters. Nat. Neurosci. 2, 375–381 (1999).

13. Carrasco, M., Penpeci-Talgar, C. & Eckstein, M. Spatial covert attention increases contrast sensitivity across the CSF: support for signal enhancement. Vis. Res. 40(10-12), 1203–1215 (2000).

14. Carrasco, M. Covert attention increases contrast sensitivity: Psychophysical, neurophysiological and neuroimaging studies. Prog. Brain Res. 154, 33–70 (2006).

15. Carrasco, M. Visual attention: The past 25 years. Vis. Res. 51(13), 1484–1525 (2011).

16. Neri, P., Parker, A. J. & Blakemore, C. Probing the human stereoscopic system with reverse correlation. Nature 401(6754), 695–698 (1999).

17. Ringach, D. & Shapley, R. Reverse correlation in neurophysiology. Cogn. Sci. 28(2), 147–166 (2004).

18. Ohzawa, I., DeAngelis, G. C. & Freeman, R. D. Stereoscopic depth discrimination in the visual cortex: neurons ideally suited as disparity detectors. Science 249(4972), 1037–1041 (1990).

19. DeAngelis, G. C., Ohzawa, I. & Freeman, R. D. Spatiotemporal organization of simple-cell receptive fields in the cat’s striate cortex. II. Linearity of temporal and spatial summation. J. Neurophysiol. 69(4), 1118–1135 (1993).

20. Nishimoto, S., Ishida, T. & Ohzawa, I. Receptive field properties of neurons in the early visual cortex revealed by local spectral reverse correlation. J. Neurosci. 26(12), 3269–3280 (2006).

21. Neri, P. & Levi, D. M. Receptive versus perceptive fields from the reverse-correlation viewpoint. Vis. Res. 46(16), 2465–2474 (2006).

22. Ringach, D. L. Tuning of orientation detectors in human vision. Vis. Res. 38(7), 963–972 (1998).

23. De Gardelle, V. & Summerfield, C. Robust averaging during perceptual judgment. Proc. Natl. Acad. Sci. 108(32), 13341–13346 (2011).

24. Hanks, T. D. & Summerfield, C. Perceptual decision making in rodents, monkeys, and humans. Neuron 93(1), 15–31 (2017).

25. Vandormael, H., Castañón, S. H., Balaguer, J., Li, V. & Summerfield, C. Robust sampling of decision information during perceptual choice. Proc. Natl. Acad. Sci. 114(10), 2771–2776 (2017).

26. Sato, H. & Motoyoshi, I. Distinct strategies for estimating the temporal average of numerical and perceptual information. Vis. Res. 174, 41–49 (2020).

27. Yashiro, R. & Motoyoshi, I. Peak-at-end rule: adaptive mechanism predicts time-dependent decision weighting. Sci. Rep. 10(1), 1–8 (2020).

28. Yashiro, R., Sato, H., Oide, T. & Motoyoshi, I. Perception and decision mechanisms involved in average estimation of spatiotemporal ensembles. Sci. Rep. 10(1), 1–10 (2020).

29. Ahumada, A. J. Jr. Perceptual classification images from Vernier acuity masked by noise. Perception 25, 2 (1996).

30. Gold, J. M., Murray, R. F., Bennett, P. J. & Sekuler, A. B. Deriving behavioural receptive fields for visually completed contours. Curr. Biol. 10(11), 663–666 (2000).

31. Eckstein, M. P., Shimozaki, S. S. & Abbey, C. K. The footprints of visual attention in the Posner cueing paradigm revealed by classification images. J. Vis. 2(1), 3–3 (2002).

32. Neri, P. & Heeger, D. J. Spatiotemporal mechanisms for detecting and identifying image features in human vision. Nat. Neurosci. 5(8), 812–816 (2002).

33. Mareschal, I., Dakin, S. C. & Bex, P. J. Dynamic properties of orientation discrimination assessed by using classification images. Proc. Natl. Acad. Sci. 103(13), 5131–5136 (2006).

34. Rajashekar, U., Bovik, A. C. & Cormack, L. K. Visual search in noise: Revealing the influence of structural cues by gaze-contingent classification image analysis. J. Vis. 6(4), 7–7 (2006).

35. Neri, P. & Levi, D. Temporal dynamics of directional selectivity in human vision. J. Vis. 8(1), 22–22 (2008).

36. Busse, L., Katzner, S., Tillmann, C. & Treue, S. Effects of attention on perceptual direction tuning curves in the human visual system. J. Vis. 8(9), 2–2 (2008).

37. Caspi, A., Beutter, B. R. & Eckstein, M. P. The time course of visual information accrual guiding eye movement decisions. Proc. Natl. Acad. Sci. 101(35), 13086–13090 (2004).

38. Okazawa, G., Sha, L., Purcell, B. A. & Kiani, R. Psychophysical reverse correlation reflects both sensory and decision-making processes. Nat. Commun. 9(1), 1–16 (2018).

39. Maruyama, H., Ueno, N. & Motoyoshi, I. Response-locked classification image analysis of perceptual decision making in contrast detection. Sci. Rep. 11(1), 1–9 (2021).

40. Faul, F., Erdfelder, E., Lang, A. G. & Buchner, A. G* Power 3: A flexible statistical power analysis program for the social, behavioral, and biomedical sciences. Behav. Res. Methods 39(2), 175–191 (2007).

41. Posner, M. I., Nissen, M. J. & Ogden, W. C. Attended and unattended processing modes: The role of set for spatial location. Modes of perceiving and processing information, 137(158), 2 (1978).

42. Carrasco, M., Ling, S. & Read, S. Attention alters appearance. Nat. Neurosci. 7(3), 308–313 (2004).

43. Ling, S. & Carrasco, M. Sustained and transient covert attention enhance the signal via different contrast response functions. Vis. Res. 46(8-9), 1210–1220 (2006).

44. Motoyoshi, I. Attentional modulation of temporal contrast sensitivity in human vision. PLoS One 6(4), e19303 (2011).

45. Ratcliff, R. A theory of memory retrieval. Psychol. Rev. 85, 59 (1978).

46. Ratcliff, R. & Smith, P. L. A Comparison of Sequential Sampling Models for Two-Choice Reaction Time. Psychol. Rev. 111, 333–367 (2004).

47. Huk, A. C. & Shadlen, M. N. Neural activity in macaque parietal cortex reflects temporal integration of visual motion signals during perceptual decision making. J. Neurosci. 25(45), 10420–10436 (2005).

48. Gold, J. I. & Shadlen, M. N. The neural basis of decision making. Annu. Rev. Neurosci. 30, 535–574 (2007).

49. Kiani, R., Hanks, T. D. & Shadlen, M. L. Bounded integration in parietal cortex underlies decisions even when viewing duration is dictated by the environment. J. Neurosci. 28(12), 3017–3029 (2008).

50. Ratcliff, R. & McKoon, G. The diffusion decision model: theory and data for two-choice decision tasks. Neural Comput. 20(4), 873–922 (2008).

51. Enroth-Cugell, C. & Robson, J. G. The contrast sensitivity of retinal ganglion cells of the cat. J. physiol. 187(3), 517–552 (1966).

52. Adelson, E. H. & Bergen, J. R. Spatiotemporal energy models for the perception of motion. Josa a 2(2), 284–299 (1985).

53. Watson, A. B. Temporal sensitivity. Handb. Percept. Human Perform. 1(6), 1–43 (1986).

54. Usher, M. & McClelland, J. L. The time course of perceptual choice: the leaky, competing accumulator model. Psychol. Rev. 108(3), 550 (2001).

55. Cheadle, S. et al. Adaptive gain control during human perceptual choice. Neuron 81(6), 1429–1441 (2014).

56. Li, V., Michael, E., Balaguer, J., Castañón, S. H. & Summerfield, C. Gain control explains the effect of distraction in human perceptual, cognitive, and economic decision making. Proc. Natl. Acad. Sci. 115(38), e8825–8834 (2018).

57. Brown, S. & Heathcote, A. A ballistic model of choice response time. Psychol. Rev. 112, 117–128 (2005).

58. Brown, S. D. & Heathcote, A. The simplest complete model of choice response time: linear ballistic accumulation. Cogn. Psychol. 57, 153–178 (2008).

